# The nature of nurture: effects of parental genotypes

**DOI:** 10.1101/219261

**Authors:** Augustine Kong, Gudmar Thorleifsson, Michael L. Frigge, Bjarni J. Vilhjálmsson, Alexander I. Young, Thorgeir E. Thorgeirsson, Stefania Benonisdottir, Asmundur Oddsson, Bjarni V. Halldórsson, Gísli Masson, Daniel F. Gudbjartsson, Agnar Helgason, Gyda Bjornsdottir, Unnur Thorsteinsdottir, Kari Stefansson

**Affiliations:** deCODE genetics/Amgen Inc., 101 Reykjavik, Iceland; Bioinformatics Research Centre, Aarhus University, 8000 Aarhus, Denmark; Department of Epidemiology, Harvard T.H. Chan School of Public Health, 02115 Boston, MA USA; Big Data Institute, Li Ka Shing Centre for Health Information and Discovery, University of Oxford, Oxford, United Kingdom; School of Engineering and Natural Sciences, University of Iceland, Reykjavik 101, Iceland; Department of Anthropology, University of Iceland, Reykjavík 101, Iceland; Faculty of Medicine, University of Iceland, Reykjavik 101, Iceland

## Abstract

Sequence variants in the parental genomes that are not transmitted to a child/proband are often ignored in genetic studies. Here we show that non-transmitted alleles can impact a child through their effects on the parents and other relatives, a phenomenon we call genetic nurture. Using results from a meta-analysis of educational attainment, the polygenic score computed for the non-transmitted alleles of 21,637 probands with at least one parent genotyped has an estimated effect on the educational attainment of the proband that is 29.9% (*P* = 1.6×10^−14^) of that of the transmitted polygenic score. Genetic nurturing effects of this polygenic score extend to other traits. Paternal and maternal polygenic scores have similar effects on educational attainment, but mothers contribute more than fathers to nutrition/heath related traits.

**One Sentence Summary:** Nurture has a genetic component, *i.e.* alleles in the parents affect the parents’ phenotypes and through that influence the outcomes of the child.

---

How the human genome (nature) and the environment (nurture) conspire to make members of our species is a fundamental question, and any novel insights there would be an important milestone. One challenge encountered by those who aspire to shed light on this is that the genome and the environment are not independent and models that fail to account for this are thus incomplete. Here we demonstrate how the genomes of close relatives, parents and siblings, can affect the proband through their contributions to the environment.

In animal studies, it is well established that alleles in a parent that are not transmitted to the offspring can nonetheless influence its phenotypes (1, 2). Most examples are on effects manifested at the fetal stage, where only the non-transmitted maternal alleles are relevant. In humans, the non-transmitted maternal alleles have been used to examine the potential causal relationships between the state of the mother during pregnancy and the outcomes of the child (3, 4). Here, for humans, we consider an alternative causal path where both paternal and maternal non-transmitted alleles can have effects that are mostly manifested after birth. A sequence variant that affects the phenotype of an individual is also likely to affect the parent from whom it was inherited (Fig. 1a). For some phenotypes, the state of a parent can influence the state of its child. This gives rise to an intriguing situation, where a child’s phenotype is influenced not only by the transmitted paternal and maternal alleles (T_P_, T_M_ in Fig. 1a), but also by the alleles that were not transmitted (NT_P_, NT_M_). A good example is educational attainment (5, 6). Thus, the educational attainment of parents provides an environmental effect for children, but one that has a genetic component (7, 8). We call this phenomenon genetic nurture. The transmitted and non-transmitted alleles (Fig. 1a) both exert effects on the parents and thus both induce genetic nurturing effects. The effect of the transmitted allele includes both its direct effect on the proband and its effect manifested through nurturing from blood relatives. Because the amount of the trait variance explained is proportional to effect size squared, genetic nurture could have a substantially bigger impact on variance explained through the transmitted alleles (by magnifying the direct effect) than the non-transmitted alleles. However, data on the non-transmitted alleles are needed to separate the genetic nurturing effects from the direct effects of the transmitted alleles. Specifically, let 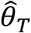 and 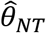 denote respectively the estimated effects of the transmitted and non-transmitted alleles when the paternal and maternal alleles are grouped together. Denoting the direct effect as δ we propose to estimate it by 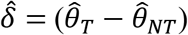. By taking the difference? genetic nurturing effects and other potential confounding effects induced by population structure and assortative mating (9, 10)(see below) are cancelled out. This approach is related to the transmission-disequilibrium test (TDT) (11, 12) as both utilize the non-transmitted alleles as controls. However, the potential effects of the non-transmitted alleles are ignored by the TDT. Mathematically, genetic nurture is a form of associative/indirect genetic effect as defined by the animal breeding literature (2). The study here, however, takes advantage of the special properties of our human data to, for the first time empirically examine the magnitudes of such effects for important traits such as educational attainment.

**Fig. 1.**
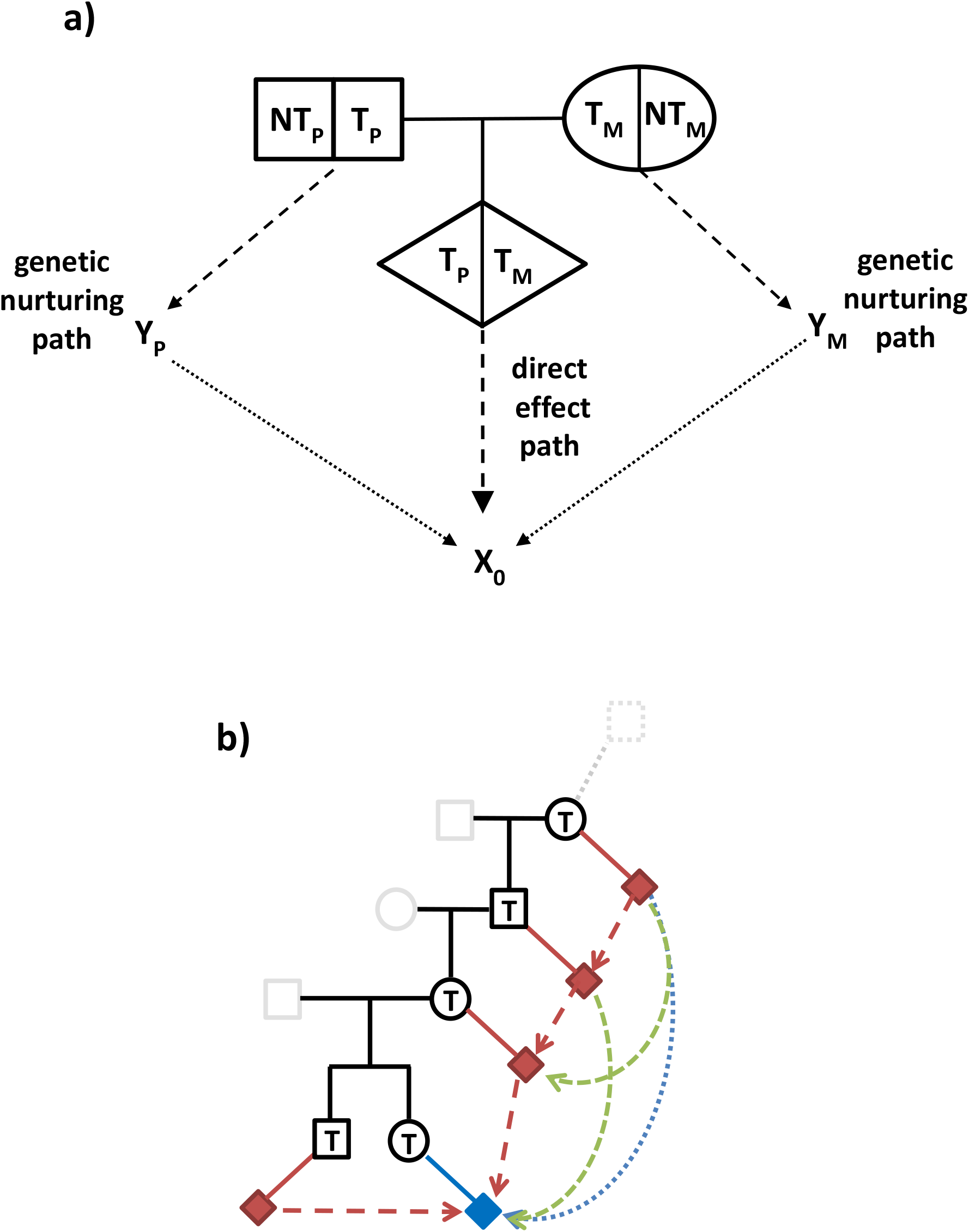
The direct genetic effect and the genetic nurturing effect. **a**, Alleles at an autosomal site carried by a parents-offspring trio are labelled with respect to the offspring/proband. T_P_ and T_M_ denote respectively the alleles transmitted from the father and the mother to the proband, and NT_P_ and NT_M_ denote the paternal and maternal alleles that are not transmitted. The transmitted alleles can influence the phenotype of the offspring, X_O_, through a direct path. The alleles of the parents, both the transmitted and the non-transmitted, can influence the parents’ phenotypes, Y_P_ and Y_M_, and through them have a nurturing effect on X_O_. This pathway combines a genetic effect, (T_P_, NT_P_, T_M_, NT_M_) on (Y_P_, Y_M_), with a nurturing effect, (Y_P_, Y_M_) on X_O_. Note that while X_O_ is often an individual trait of interest, Y would include a much broader set of phenotypes and not completely known. **b**, Red diamonds denote phenotypes of relatives. Blue diamond denotes phenotype of the proband. Using the maternally transmitted allele as an example (denoted by T), we highlight that, in addition to the parents, the genetic nurturing effect can be manifested through the phenotypes of older ancestors and non-ancestors such as sibling.

## Estimating direct effects

To maximize the power to detect the effects of the non-transmitted alleles, we constructed polygenic scores (13) using 618,762 single nucleotide polymorphisms (SNPs) spanning the genome. The per-locus allele-specific weightings for the polygenic scores were derived from applying LDpred (14) to the results of a large genomewide association study (8) (GWAS) of educational attainment measured in years of education (EA), with Icelandic data removed (see Supplementary Material). The first analysis focused on 21,637 Icelandic probands with EA data, born between 1940 and 1983 (9,139 males, 12,498 females), and with at least one parent genotyped (Table 1). Since we could determine the parent of origin of the transmitted alleles (15), the non-transmitted allele from a genotyped parent is easily determined. Let poly_TP_ and poly_TM_ be the polygenic score computed from the transmitted paternal and maternal alleles respectively, and let poly_NTP_ and poly_NTM_ denote the corresponding polygenic scores for the non-transmitted alleles. To maximize power, we start by providing the results for poly_T_ = poly_TP_ + poly_TM_ and poly_NT_ = poly_NTP_ + poly_NTM_, Here, poly_TP_ and poly_TM_ are scaled so that poly_T_ have mean zero and variance 1, and the trait EA is also standardized to have variance 1. Poly_NTP_ and poly_NTM_ were similarly computed, and a zero was imputed when the parent was not genotyped (see Supplementary Material). Associations between EA and the polygenic scores computed based on a joint analysis of poly_T_ and poly_NT_ that adjusts for sex, year of birth (yob) up to the cubic term, interactions between sex and yob, and 100 principal components (PCs) (see Supplementary Material) are presented in Table 1. The estimated effect of poly_T_, 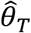, is 0.223 and highly significant (*P* = 1.6 ×10^−174^, calculated with genomic control adjustment (see Supplementary Material)(16)). Since both poly_T_ and EA are standardized, the estimated fraction of the trait variance explained by poly_T_ is 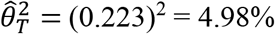 (*R*^2^ in Table 1). However, the estimated effect of poly_NT_, 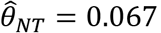, is also highly significant (*P* = 1.6 ×10^−14^). Thus the estimated direct effect of poly_T_, 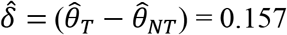, only explains 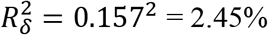 of the trait variance, approximately one-half of *R*^2^. Noting that 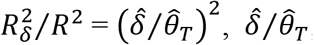, *i.e.* 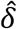 as a fraction of 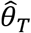, is presented in Table 1. In addition to the polygenic scores, individual results for 120 SNPs that are genomewide significant (*P* < 5 ×10^−8^) in the Iceland-excluded meta-analysis are provided (Supplementary Table S1). Fifteen of the 120 SNPs (12.5%) have one-sided *P* < 0.05 for the non-transmitted alleles, more than what are expected from noise (*P* = 1.5 ×10^−3^, (see Supplmentary Material)). Results here are consistent with previous observations that within-family effects calculated using dizygotic twins for EA were overall smaller than the standard GWAS effect estimates (7, 8).

**Table 1.** Decomposition of the observed effect of the polygenic score into direct, genetic nurturing, and confounding effects

|  |  |  |  | Transmitted |  |  | Nontransmitted |  |  |  |  |  |  |
| --- | --- | --- | --- | --- | --- | --- | --- | --- | --- | --- | --- | --- | --- |
|  |  |  |  | T (T = T<sub>P</sub> + T<sub>M</sub>) |  |  | NT (NT = NT<sub>P</sub> + NT<sub>M</sub>) |  |  |  |  |  |  |
| Trait | N | N<sub>NTP</sub> | N<sub>NTM</sub> | $\hat{\theta}_T$ | P | R<sup>2</sup> (%) | $\hat{\theta}_{NT}$ | P | R<sub>δ</sub><sup>2</sup> (%) | $\hat{\delta} / \hat{\theta}_T$ | $\hat{\phi}_\delta / \hat{\theta}_T$ | $\hat{\eta} / \hat{\theta}_T$ | $\hat{\phi}_\eta / \hat{\theta}_T$ |
| EA | 21637 | 13948 | 19012 | 0.223 | 1.6×10<sup>-174</sup> | 4.98 | 0.067 | 1.6×10<sup>-14</sup> | 2.45 | 0.701 | 0.046 | 0.224 | 0.029 |
| AGFC | 54372 | 35294 | 47052 | 0.108 | 9.7×10<sup>-110</sup> | 1.17 | 0.039 | 2.9×10<sup>-13</sup> | 0.48 | 0.640 | 0.052 | 0.264 | 0.043 |
| HDL | 46872 | 30855 | 40788 | 0.065 | 9.0×10<sup>-29</sup> | 0.42 | 0.027 | 6.0×10<sup>-6</sup> | 0.14 | 0.586 | 0.046 | 0.319 | 0.050 |
| BMI | 39078 | 26433 | 34533 | -0.060 | 1.0×10<sup>-22</sup> | 0.36 | -0.017 | 0.0077 | 0.19 | 0.718 | 0.055 | 0.197 | 0.030 |
| FG | 34767 | 22959 | 30222 | -0.051 | 7.6×10<sup>-18</sup> | 0.26 | -0.018 | 0.0059 | 0.11 | 0.655 | 0.052 | 0.252 | 0.040 |
| HT | 39270 | 26563 | 34703 | 0.052 | 6.6×10<sup>-14</sup> | 0.28 | 0.030 | 1.5×10<sup>-5</sup> | 0.05 | 0.422 | 0.031 | 0.476 | 0.071 |
| CPD | 18887 | 12371 | 16589 | -0.055 | 1.4×10<sup>-12</sup> | 0.31 | -0.030 | 5.3×10<sup>-4</sup> | 0.06 | 0.461 | 0.035 | 0.439 | 0.066 |
| HLTH | 62328 | 41996 | 54546 | 0.082 | 2.7×10<sup>-60</sup> | 0.67 | 0.033 | 8.9×10<sup>-11</sup> | 0.23 | 0.592 | 0.051 | 0.305 | 0.052 |

**Trait:** educational attainment (EA), age at first child (AGFC), high-density lipoprotein level (HDL), body mass index (BMI), fasting glucose level (FG), height (HT), cigarettes per day for smokers (CPD), composite health trait (HLTH). Traits are standardized to have variance one. **N:** number of probands with at least one parent genotyped; **N<sub>NTP</sub>:** number with father genotyped; **N<sub>NTM</sub>:** number with mother genotyped. $\hat{\theta}_T, \hat{\theta}_{NT}$ : estimated effects of the polygenic scores computed for the transmitted and non-transmitted alleles when they are analysed jointly. $\hat{\delta} = (\hat{\theta}_T - \hat{\theta}_{NT})$ : estimated direct effect of the polygenic score. **R<sup>2</sup>:** estimated variance accounted for by the transmitted polygenic score, which captures both the direct effect and the genetic nurturing effect. **R<sub>δ</sub><sup>2</sup>:** estimated variance accounted for by the direct effect alone. $\hat{\phi}_\delta, \hat{\eta}, \hat{\phi}_\eta$ : estimate, respectively, the assortative mating induced confounding effect for the direct effect component, the genetic nurturing effect, and the confounding effect of the genetic nurturing component.

## Assortative mating and estimating the genetic nurturing effect

Let *η* denote the magnitude of the genetic nurturing effect. Even though our analyses have adjustment for 100 PCs, which should have eliminated much of the population stratification induced confounding, 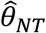 can still be capturing effects other than *η*. When there is assortative mating with respect to the genetic component underlying EA (10), a subtle confounding effect can result. Fig. 2 illustrates a simple scenario where the phenotype is assumed to be influenced by two loci A and B. If there is assortative mating in the parents’ generation, it would lead to correlation of alleles between partners, e.g. the A alleles of the father (A_1_ and A_2_ in Fig. 2) will be correlated with the B alleles of the mother (B_3_ and B_4_), and vice versa. Consequently, the paternally transmitted A allele A_P_ will be positively correlated with the maternally transmitted B allele B_M_, and A_M_ will be correlated with B_P_. This correlation between alleles inherited from different parents is referred to as *trans* correlation, while the correlation between alleles inherited from the same parent, e.g. A_P_ and B_P_, is referred to as *cis* correlation. This assortative mating induced correlation differs from correlation between markers that are close physically, *i.e.* within the same linkage-disequilibrium block. The latter correlation is mainly driven by the *cis* component, while the assortative mating induced correlation could be dominated by the *trans* component. If trait association is calculated for locus A individually, the observed effect will capture both the effect of locus A and part of the effect of B. Let *ϕ_δ_* denote this added confounding effect. Similarly, assortative mating would also lead the A alleles to capture some of the nurturing effect of B, an effect denoted by *ϕ_η_*. Under our model assumptions (see Supplementary Material),

**Fig. 2.**
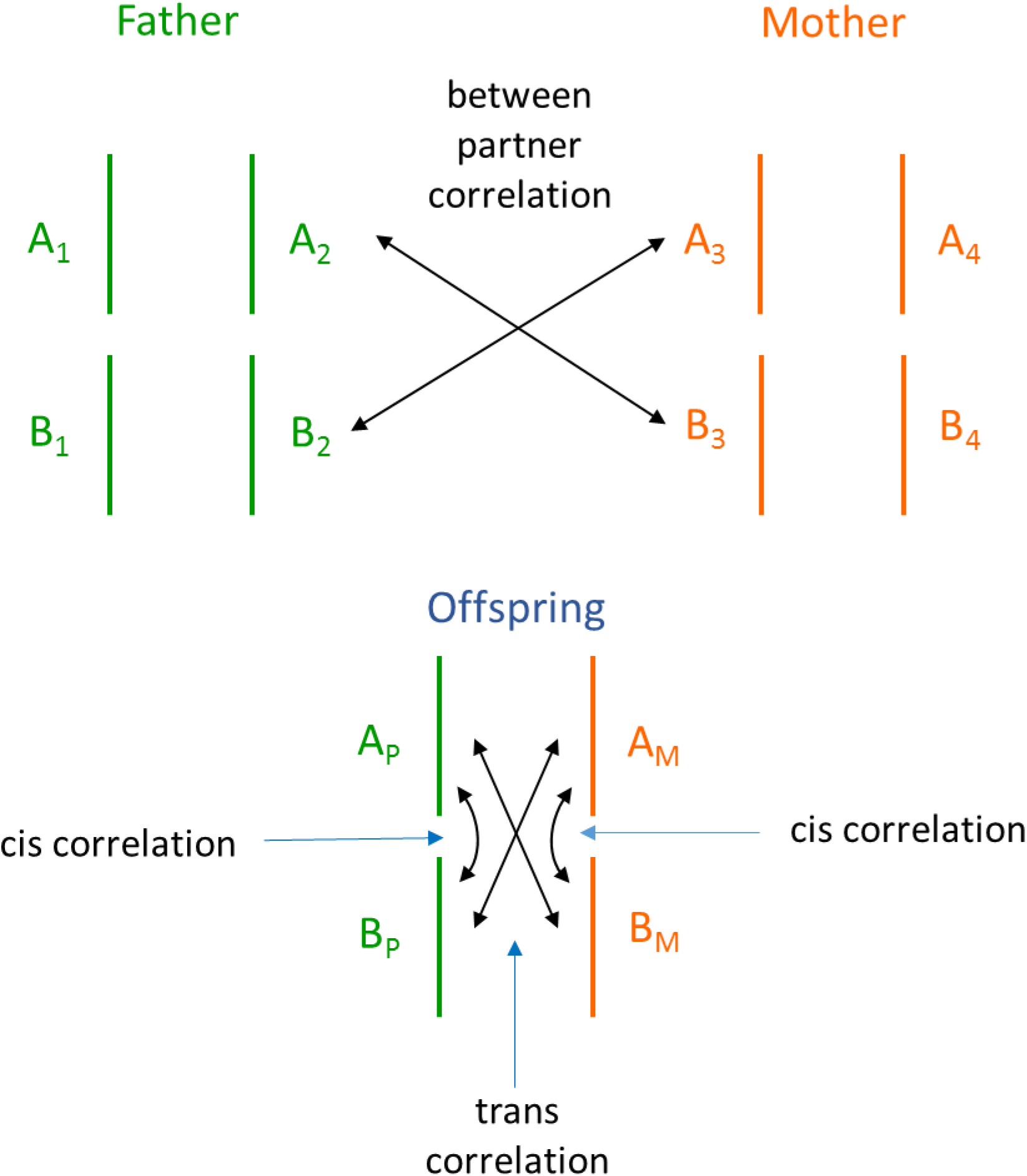
Correlation and confounding induced by assortative mating. An example of two loci, A and B, contributing to the phenotype. Through assortative mating, alleles in the father become correlated with alleles in the mother. Consequently, the transmitted paternal alleles (A_P_, B_P_) become correlated with the maternally transmitted alleles (A_M_, B_M_). This correlation between alleles with different parental origins is referred to as *trans* correlation, while correlation between alleles with the same parental origins, *e.g*. A_P_ and B_P_, is referred to as *cis* correlation. When A_P_/A_M_ and B_M_/B_P_ are correlated, association analysis between the phenotype and A alone will also capture part of the effect of B.

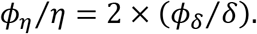

The factor of 2 arises because the non-transmitted alleles have the same nurturing effects as the transmitted ones, and thus the transmitted/non-transmitted A alleles are capturing, through correlation, the nurturing effects of both the transmitted and non-transmitted B alleles. We have the decompositions

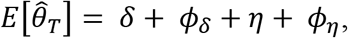

and

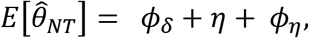

where *E*[] denotes expectation. Because both the transmitted and non-transmitted A alleles capture the confounding effects, 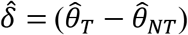 remains an appropriate estimate of the direct effect *δ*. Locus A and locus B in Fig. 2 can be generalized to represent two non-overlapping sets of loci. For the study here, we think of A as the EA polygenic score, while B represents the genetic component of EA that is statistically orthogonal to A (under a scenario of no assortative mating). The mathematical relationships highlighted above continue to hold for the polygenic scores either exactly or approximately. Using a novel method for estimating heritability that also utilizes data on the non-transmitted alleles (17), the full genetic component of EA is estimated to have a direct effect that explains 17.0% of the variance of EA. In other words, poly_T_ is estimated to be 2.45/17.0 = 14.4% of the full genetic component, while the remaining 85.6% corresponds to the B components. Based on this estimate, we extrapolate the correlations observed between the paternal polygenic scores (poly_TP_ and poly_NTP_) and the maternal polygenic scores (poly_TM_ and poly_NTM_) to estimate the correlations between them and the unobserved B components (see Supplementary Material). From the latter, *ϕ_δ_*/*δ* and *ϕ_η_*/*η* are estimated as 0.065 and 0.130 respectively. This estimation avoided making the assumption that assortative mating between parents was manifested only through correlation of their educational attainments, which would have led to lower estimates for the *ϕ*’s (see Supplementary Material). From these estimates and the above equations, 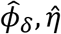, and 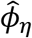 were computed and presented in Table 1 as fractions of 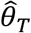. For EA, 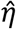 accounts for about 75% of the value of 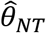 and is 31.9% the size of 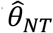. Finally, we note that assortative mating occurring before the parents’ generation could lead to additional confounding. However, this effect appears negligible here as, after adjustment for 100 PCs, the within-parent correlation of the transmitted and non-transmitted polygenic scores is actually negative (but *P* > 0.05) (see Supplementary Material).

## Direct and nurturing effects on other traits

The educational attainment polygenic score have significant associations with many other quantitative traits in our database. Among them, those with the strongest statistical significance are age-at-first-child (AGFC) (18), high-density lipoprotein (19) (HDL), body mass index (20) (BMI), fasting glucose level (21) (FG), height (22) (HT), and cigarettes smoked per day by smokers (23) (CPD). The effects of the transmitted and non-transmitted EA polygenic scores on these phenotypes were estimated as before for the EA phenotype (Table 1). For all of these traits, even though the fraction of variance explained by poly_T_ (*R*^2^) is smaller than that for EA, the effect of poly_NT_ is statistically significant. Moreover, except for BMI, the ratio 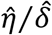, is higher for these traits than for EA, and exceeds 1 for HT.

## Parent of origin

Table 2 provides the estimated effects of poly_TP_, poly_TM_, poly_NTP_ and poly_NTM_ separately (see Supplementary Material). For EA, 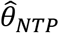, the estimated effect of poly_NTP_, is highly significant (*P* = 5.2 ×10^−7^), and its value is nearly identical to 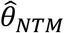 (higher *P* for poly_NTP_ is due to fewer fathers genotyped than mothers). This indicates that the effect previously observed for poly_NT_ is not driven by epigenetic effects such as imprinting or genetic interactions between foetus and mother in the womb, and is indeed capturing a genetic nurturing effect (also see Supplementary Tables S2 and S3 which have results for polygenic scores calculated without SNPs in imprinted regions(24)). However, even with both parents contributing to genetic nurture, the magnitude of the effect can differ between fathers and mothers. Let *η_P_* and *η_M_* denote respectively the paternal and maternal genetic nurturing effect. Since the transmitted alleles also contribute to the nurturing effect, we use a weighted average of 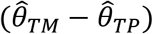 and 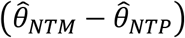, with weights proportional to (standard error)^−2^ (see Supplementary Material), to estimate (*η_M_* – *η_P_*) (Table 2). Combining this estimate with 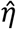 from Table 1, considered as an estimate of a weighted average of *η_P_* and *η_M_* with weights proportional to the numbers of fathers and mothers genotyped, we calculated individual estimates of *η_P_* and *η_M_* (see Supplementary Material), denoted by 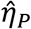 and 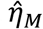, and the ratio 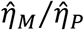 (Table 2). For EA, (*η_M_* – *η_P_*) is estimated to be 0.011, but it is not significantly different from zero (*P* = 0.31), *i.e.* the ratio 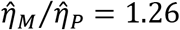 is not significantly different from 1. For all of the other six traits, 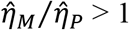, but nominally significant only for HT 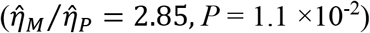. HDL and FG have *P* between 0.05 and 0.10. To increase power, for individuals for whom we had data for one or more of the five health/nutrition related traits (HDL, BMI, FG, HT, and CPD), a composite health trait (HLTH) was constructed by taking the sum of the standardized values of the available traits (positive signs for HDL and HT, and negative signs for BMI, FG and CPD) and dividing it by the square-root of the number of trait values summed. It was then standardized to have variance one. For HLTH, 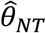 has a larger value than that for the individual health/nutrition trait and highly significant (*P* = 8.9 ×10^−11^, Table 1). Both 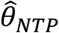 and 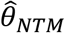 are significant, but 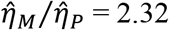 with a *P* of 4.8 ×10^−3^ (Table 2). This supports the notion that mothers have a stronger nurturing effect than fathers do on the health of the child.

**Table 2.** Parent-of-origin specific effects of the polygenic scores

| Trait | Transmitted | | | | Non-Transmitted | | | | | | $\hat{\eta}_M / \hat{\eta}_P$ |
| --- | --- | --- | --- | --- | --- | --- | --- | --- | --- | --- | --- |
| | $\hat{\theta}_{TP}$ | $P$ | $\hat{\theta}_{TM}$ | $P$ | $\hat{\theta}_{NTP}$ | $P$ | $\hat{\theta}_{NTM}$ | $P$ | Estimate of $\eta_M - \eta_P$ | $P$ | |
| EA | 0.214 | $1.0 \times 10^{-89}$ | 0.232 | $9.9 \times 10^{-103}$ | 0.066 | $5.2 \times 10^{-7}$ | 0.067 | $3.6 \times 10^{-9}$ | 0.011 | 0.31 | 1.26 |
| AGFC | 0.100 | $1.4 \times 10^{-52}$ | 0.116 | $1.9 \times 10^{-68}$ | 0.034 | $2.7 \times 10^{-5}$ | 0.043 | $1.2 \times 10^{-9}$ | 0.013 | 0.067 | 1.59 |
| HDL | 0.062 | $5.0 \times 10^{-16}$ | 0.068 | $7.8 \times 10^{-19}$ | 0.013 | 0.13 | 0.037 | $2.0 \times 10^{-6}$ | 0.014 | 0.077 | 2.05 |
| BMI | -0.059 | $5.2 \times 10^{-13}$ | -0.062 | $1.2 \times 10^{-13}$ | -0.019 | 0.055 | -0.016 | 0.062 | -0.000 | 0.98 | 1.02 |
| FG | -0.043 | $1.0 \times 10^{-7}$ | -0.059 | $4.1 \times 10^{-13}$ | -0.011 | 0.27 | -0.023 | 0.0073 | -0.014 | 0.090 | 3.99 |
| HT | 0.035 | $8.8 \times 10^{-5}$ | 0.070 | $1.3 \times 10^{-14}$ | 0.027 | 0.0082 | 0.033 | $3.5 \times 10^{-4}$ | 0.023 | 0.011 | 2.85 |
| CPD | -0.042 | $1.3 \times 10^{-4}$ | -0.069 | $2.6 \times 10^{-10}$ | -0.036 | 0.0071 | -0.025 | 0.028 | -0.011 | 0.33 | 1.63 |
| HLTH | 0.070 | $1.4 \times 10^{-26}$ | 0.093 | $1.0 \times 10^{-44}$ | 0.026 | $7.5 \times 10^{-4}$ | 0.039 | $8.4 \times 10^{-9}$ | 0.019 | 0.0048 | 2.32 |

## Variance explained and effects of siblings

The existence of genetic nurture complicates the estimation and interpretation of heritability (17), which has been recognized in the animal breeding literature for maternal effects (25). While distinct from a direct effect, genetic nurture is nonetheless a real effect. Notably, if there are two uncorrelated variants of the same frequency, one having a direct effect *δ* only, and the other having a nurturing effect *η* only, then the variance explained is proportional to *δ*^2^ + *η*^2^. By comparison, if one variant has both effects, then the variance explained is proportional to (*δ* + *η*)^2^ = *δ*^2^ + 2*δη* + *η*^2^ (Fig. 3), with the extra 2*δη* term. Moreover, (*δ* + *η*)^2^ only captures the effect of the transmitted allele(s), the phenotypic variance accounted for by the transmitted and non-transmitted alleles together is proportional to (*δ* + *η*)^2^ + *η*^2^ (Fig. 3). With 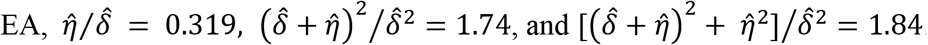. Assuming that the direct effect alone explains 17.0% of the variance, the variance explained by the transmitted alleles with the nurturing effects included would be 17.0% × 1.74 = 29.6%. Including also the non-transmitted alleles would increase variance explained to 17.0% × 1.84 = 31.3%. The genetic nurturing effect not only magnifies the variance explained, it induces an even larger amplification of the phenotypic correlations of parents and offspring and of siblings (Fig. 3, (see Supplementary Material)). Also worth noting is that the 2*δη* term highlighted above does not exist for adopted children, as then both alleles of a parent would be non-transmitted.

**Fig. 3.**
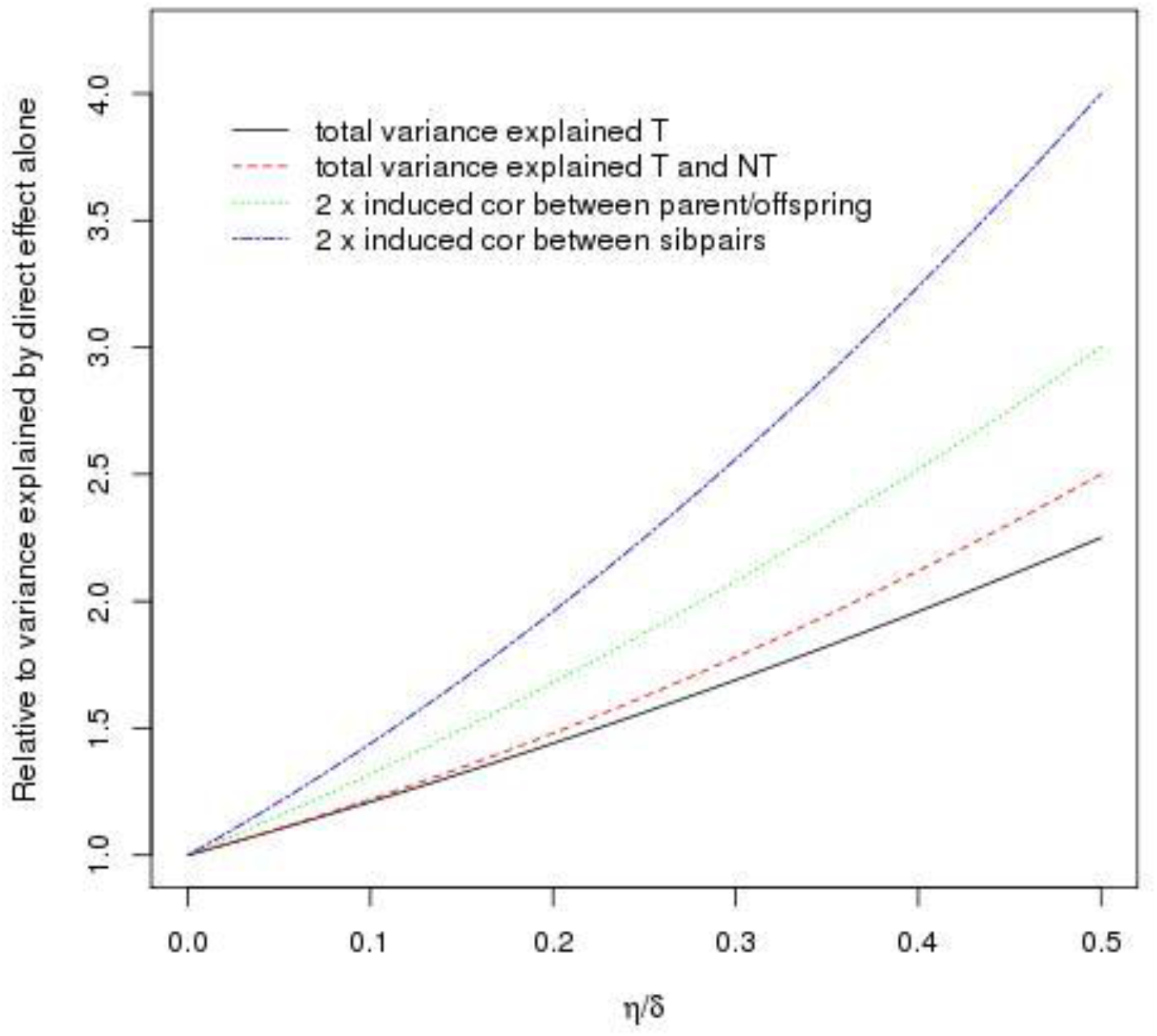
Variance explained and induced correlations between phenotypes of parent-offspring and siblings. Results are displayed as a function of the ratio *η/δ*. The y-axis is the relative amplification, *i.e.* various measures relative to what can be accounted for by the direct effect alone, the latter proportional to *η*^2^. The total variance explained by the transmitted alleles is proportional to (*δ* + *η*)^2^ (the plotted curve is hence (*δ* + *η*)^2^/*δ*^2^), while the total variance explained by the transmitted alleles plus the non-transmitted alleles is proportional to (*δ* + *η*)^2^ + *η*^2^. Formulas for the induced parent-offspring and sibling correlations are derived and given in Supplementary Material.

When introducing genetic nurture, the effect manifested through the phenotypes of the parents was emphasized. However, there can be additional contributions, although probably substantially smaller, going through grandparents and great grandparents, etc. (Fig. 2b). Furthermore, if the phenotype of the proband can be directly influenced by the phenotypes/behaviour of a sibling, as proposed in a recent paper (26), then part of genetic nurture could go through a sibling. Based on the genealogy, for each EA proband who has at least one sibling, the sibling most likely to have the biggest effect on the proband was identified as follows. If the proband has older siblings, the older sibling with yob closest to the proband was selected (monozygotic twins were excluded, but we count a dizygotic twin of the proband as an older sib). If the proband is the eldest child, a younger sibling with the closest yob was chosen. There are 7,798 probands whose chosen sibling is genotyped and whose parents are both genotyped. A polygenic score, denoted by poly_TS_, was computed using the alleles transmitted from the parents to the sibling. The educational attainment of the proband was then regressed on poly_T_, poly_NT_ and poly_TS_ jointly. The effect of poly_TS_ is significant (*P* = 0.015) and is estimated to be 24.1% (95% confidence interval: 4.7% to 43.6%) of the direct effect. The uncertainty is large because poly_TS_ is strongly correlated with poly_T_ and poly_NT_. One compensation is that, having adjusted for both poly_T_ and poly_NT_, the estimated effect of poly_TS_ is free of confounding from assortative mating.

Heritability is defined as the fraction of phenotypic variance explained by direct effects alone. The presence of parental genetic nurture introduces bias to estimates of heritability from GREML-type methods (27), such as embodied in the software package *GCTA* (28), that use correlations due to transmitted alleles without distinction between direct genetic effects and genetic nurturing effects (17). By contrast, heritability estimates based on comparing correlations between monozygotic versus dizygotic twins (29) are unaffected as the effects of parental genetic nurture are cancelled out. However, when genetic nurturing effects that go through the phenotypes of a sibling/twin are present, then this would affect both twin-based heritability estimates (30) and estimates from GREML-type methods.

## The nature of genetic nurture and other polygenic scores

To utilize the EA trait data we have for many parents, we performed analyses that treated the non-transmitted polygenic score of a genotyped parent as missing if the EA of that parent was unknown. For these data, (unadjusted) estimates of *θ_NT_* were calculated as before (Supplementary Table S4). Also given are estimates of *θ_NT_* adjusted for the EAs of the parents, obtained by adding the latter to the explanatory variables in the regressions. Notably, for EA, AGFC, HT, and HLTH, the adjusted estimate remains highly significant (*P* < 0.005), and the ratio of the adjusted versus unadjusted estimate is, respectively, 47.6%, 63.0%, 80.3%, and 68.6%. This indicates that the EA of the parent is indeed an important part of the parental phenotypes (Y in Fig. 1a) through which genetic nurture operates, but it is far from all of it.

To contrast the results presented for the polygenic score constructed from a GWAS of EA (EA polygenic score), we examined polygenic scores constructed from GWASs of height (31) (HT polygenic score) and BMI (32) (BMI polygenic score). Results corresponding to Table 1 are in Supplementary Tables S5 and S6. Noting that the HT and BMI polygenic scores are, respectively, positively (*r* = 0.087) and negatively correlated (*r* = −0.146) with the EA polygenic score, we computed HT and BMI polygenic scores adjusted for the EA polygenic score by regressing the former on the latter and taking the residuals. Their estimated effects are in Supplementary Tables S7 and S8. While the non-transmitted polygenic score has a few significant associations in Supplementary Tables S5 and S6, in Supplementary Tables S7 and S8, the only significant effect of the non-transmitted polygenic score is between the HT trait and the non-transmitted HT polygenic score. Furthermore, most of this observed effect is estimated to be due to assortative mating confounding.

## Discussion

We introduced the concept of genetic nurture and through the study of the non-transmitted alleles demonstrated that genetic nurturing effects exist, and can have a substantial impact on variance explained. These results also revealed that the observed effects from GWAS are not necessarily reflecting the direct effects alone. They can be amplified by genetic nurturing effects and to a lesser extent, assortative mating induced confounding. Due to power considerations, we mostly studied variants as an aggregate. It is however clear, given the complexity of the educational attainment trait (6) and our observed effects of the EA polygenic score on other traits, that for individual variants, the ratio of the genetic nurturing effect versus the direct effect must have variations both between and within traits. Given enough data, analyses incorporating the non-transmitted alleles would add insight into the pathway(s) through which the effect of an individual variant is manifested, as well as a better understanding of some pleiotropic effects (33).

While it is not a novel concept that genes can affect the environment (23, 34, 35), the contribution of a genetic effect manifested through nurturing has mostly been ignored in GWAS. Results here highlight the importance of family data.

The focus has been on genetic nurture in one direction, but the effects are reckoned to be in general bidirectional. For a parent-offspring pair, the magnitude of the effect in the direction of parent to offspring is likely to dominate the effect in the opposite direction. However, with siblings/twins, the effects would be reciprocal.

Analyses here implicitly assumed that direct genetic effects and genetic nurturing effects are additive, but interactive effects could certainly exist. Moreover, alleles other than the four in the parents can also have an effect, e.g. the genetic makeup of the population where the proband grew up in could be an important environmental contributor to his phenotypes.

## Acknowledgments

We thank Aysu Okbay for providing the EA meta-analysis results with Icelandic data removed, and Joel Hirshhorn for pointing out that the non-transmitted alleles could be capturing some confounding effects due to assortative mating. We thank the GIANT consortium, Andrew Wood, and Adam Locke for assisting us to obtain meta-analysis results for height and BMI with Icelandic data removed. A summary of the data that have been utilized for this manuscript is in Supplementary Table S9.

